# ABCP_finder: A Transformer Embedding-Based Prediction of Anti-Breast Cancer Peptides

**DOI:** 10.64898/2026.09.06.749767

**Authors:** Prabhat Tripathi, Rahul Semwal, Ankish Arya, Nirmalya Sen, Pritish Kumar Varadwaj

**Affiliations:** Department of Applied Sciences, Indian Institute of Information Technology Allahabad, Prayagraj, India; Department of Computer Science & Engineering, Indian Institute of Information Technology Nagpur, India; Department of Biological Sciences, Bose Institute, West Bengal, India

**Keywords:** artificial intelligence, drug discovery, anticancer peptides, breast cancer, precision medicine

## Abstract

Breast cancer remains one of the leading causes of cancer-related deaths among women worldwide. Anti-breast cancer peptides (ABCPs) have emerged as effective drug candidate due to its low toxicity, high selectivity, and ability to target cancer. However, it is very time consuming and expensive to identify novel ABCPs only through experiment methods. To address this challenge, we developed ABCP_finder, the first dedicated computational framework specifically designed for the prediction of ABCPs. The curated dataset comprised of 301 experimentally validated ABCPs and 1052 non-ABCPs. Unlike general anticancer peptide (ACP) predictors which are not trained on breast cancer specific datasets. It was trained using deliberately stringent and biologically relevant ABCPs and non-ABCPs. Instead of random train-test split, homology-aware split using CD-HIT was performed to prevent data leakage. Peptide representations generated using ProtBERT and ESM2 were classified using a multilayer perceptron (MLP). Among the tested models, ProtBERT-based model showed superior performance, achieving 93.82% accuracy, 86.88% recall, 90.59% F1-score, 0.8618 MCC, 96.67% AUC, and a Brier Score of 0.0633. Calibration analysis supported the selection of a 0.7 probability threshold for identifying high confidence ABCPs. Further external validation using xDeep-AcPEP demonstrated that peptide predicted as ABCPs by ABCP_finder are exhibiting favourable IC□□ values. ABCP_finder provides a reliable platform for high throughput large-scale ABCP screening of novel peptide therapeutics for breast cancer treatment.

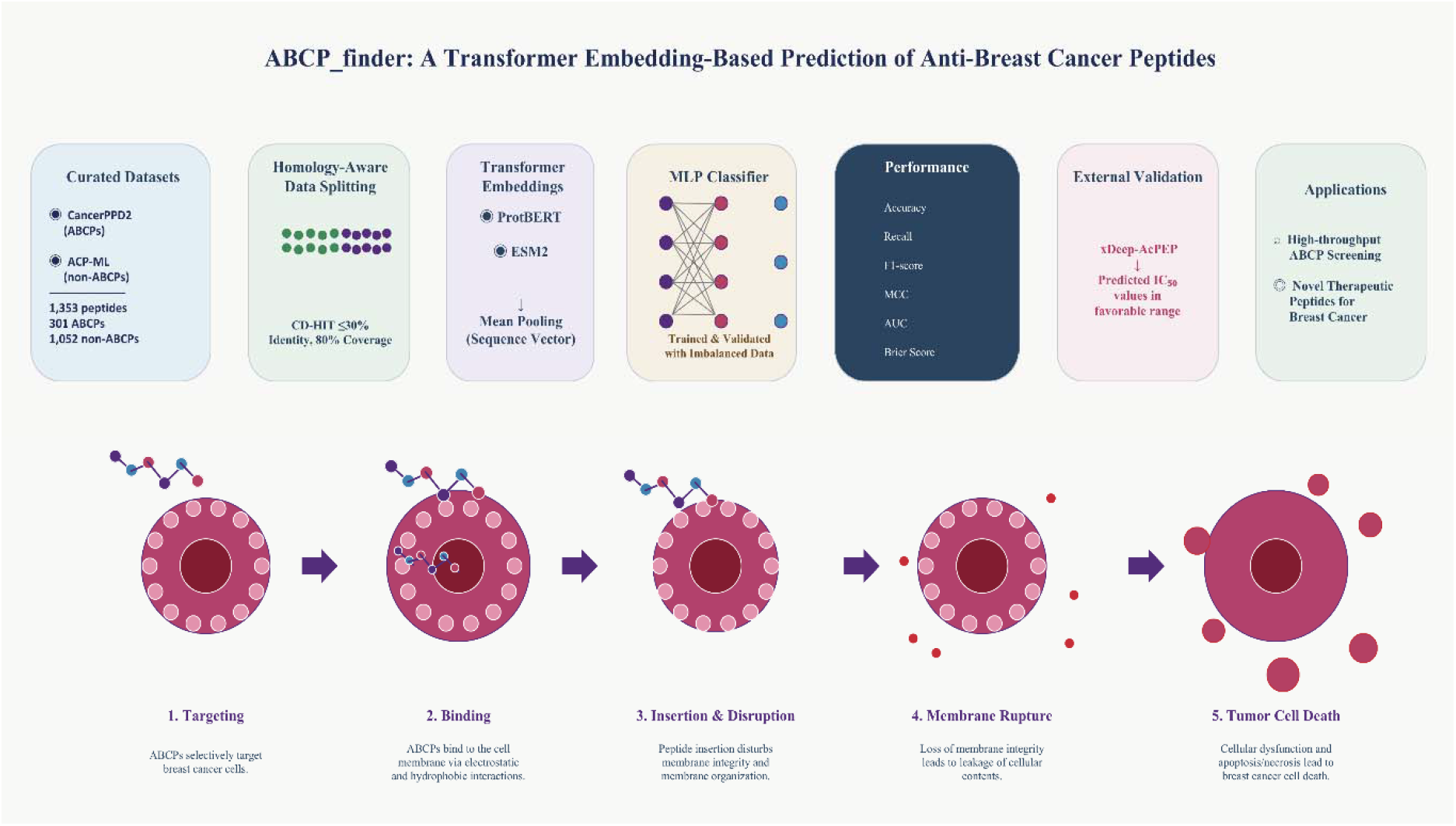

Caption: Figure illustrating the graphical abstract of ABCP_finder.

## 1. Introduction

Breast cancer is one of the most common and lethal diseases among women globally [1]. Conventional therapies are facing difficulties due to aggressiveness and drug resistance of tumours [2]. Although these conventional methods have improved the health of patients, after long usage the efficacy of these treatment options is limited by systemic toxicity, non-target effects, and drug resistance [2]. To address these limitations, peptides have come up as a promising alternative because of their high selectivity, biocompatibility, and comparatively negligible immunogenicity [3]. For example, LYP-1 (CGQKRTRGC) targets the mitochondrial p32 protein of melanoma cancer. Another peptide, IL4RPep-1 (CRKRLDRNC), has been a target of the interleukin-4 receptor and (KLAKLAK)□ by triggering apoptosis and found associated with therapeutic applications in breast, lung, and colon cancers [4].

Peptide-based treatments in breast cancer can precisely target overexpressed receptors, cellular proteins, and transcription factors such as EGF, BCL2, HER2, EGFR, and particularly cell membrane proteins [5-7]. Anti-Breast Cancer Peptides (ABCPs) are short peptides (<50 amino acids) that have emerged as an efficient and advantageous candidate to inhibit breast cancer through multiple mechanisms of action, for example, triggering apoptosis; destroying cell membranes; inhibiting signalling pathways; and inducing immunological responses while minimizing adverse effects on healthy cells [8, 9]. Several experimental peptides have shown promising activity towards breast cancer. Lactoferricin B, sourced from bovine lactoferrin; KLA ([KLAKLAK]□); P18, derived from *Arhgdia;* Aurein and Citropin (derived from frog skin) and Melittin (derived from bee venom) represent promising candidates for the development of potential and novel anticancer peptides (ACP) towards breast cancer [10-14].

In addition to these experimental ACP, A Food and Drug Administration approved peptide, Goserelin, which is a Gonadotropin releasing hormone analogue, slows down hormone dependent breast cancer by inhibiting ovarian oestrogen production in post-menopausal women [15].Together, these attributes create ABCPs a promising and highly specific strategy for breast cancer treatment, highlighting the need of the development of novel ABCPs.

Traditional computational methods for peptide prediction are mainly relied on physicochemical and evolutionary features derived from peptide sequences, such as amino acid composition, dipeptide composition, hydrophobicity profiles, BLOSSUM62, charge, and other biochemical features [16-18]. These basic features are then used as inputs for machine learning algorithms to separate out peptides into different classes based on their biological activities. Although, those approaches were efficient in terms of computation cost and easily interpretable, but they often get failed to capture the complex sequence patterns, structural characteristics and interactions that plays very important role in determining peptide activity. Because these features only provide simplified representations of peptide sequences. For example, amino acid, dipeptide-, and tripeptide-composition features summarize the sequence content. These features can not distinguish two peptides with identical composition but different residue order. This makes the resulting model may struggle to learn deeper biological relationships within the data. Ultimately, the ability of accurate prediction of activity of novel peptides and generalisation to unknow peptides remains restricted. So, this kind of long range and order dependent information is precisely captured with context aware embeddings.

The integration of deep learning and transformer based pretrained models has significantly increased the advancement of sequence-based prediction task. This integration enables the more accurate and biologically meaningful analysis compared to traditional approaches [19]. Models like ESM2 (Evolutionary Scale model 2) [20] and ProtBERT (Protein Bidirectional Encoders Representations from Transformers) [21], which draw inspiration from natural language processing (NLP) interpret amino acids as tokens inside a protein sequence. By examining the sequential context of these tokens, model understands complicated relationships within protein sequences and generates high-dimensional embeddings that capture both local and global interactions among neighbouring residues throughout the whole sequence. These embeddings intrinsically encode crucial structural, biochemical, and evolutionary information, removing the need of manual and labour-intensive feature engineering while offering deeper illustration of peptide sequences [22].

ProtBERT is a protein language model built on BERT architecture, originally designed for natural language processing tasks. It was developed as the part of ProtTrans initiative with the goal of understanding protein sequences in a similar way that language models understand sentences. ProtBERT is pretrained on massive sequence databases such as Big Fantastic Database (BFD) [23, 24]and UniRef [25], which contains billions of amino acid sequences. During training it utilizes masked language modelling approach where amino acid sequences are randomly hidden and the model learn to predict them on the basis of surrounding contexts.

The self-supervised learning strategy enables ProtBERT to capture long range interactions between amino acids, evolutionary patterns across families, functional motifs and sequence signatures. These characteristics makes ProtBERT to generate rich numerical representations of protein sequences, which can be effectively used for down streaming tasks such as classification, functional prediction drug discovery applications.

ESM2 is a state-of-the-art protein language model developed to learn directly from raw sequences at an unprecedented scale. Similar to ProtBERT, it also uses the masked language modelling objective, but it is trained on even larger and diverse datasets. One key advantage is it do not rely on multiple sequence alignment to capture evolutionary patterns. ESM2 learns evolutionary patterns implicitly from sequence data alone. Using this training approach, ESM2 is able to capture fine-grained residue level interactions, understands global structural and functional relationships and generalise across diverse protein families. Because of these capability ESM2 can be used in highly bioinformatics tasks such as protein structure prediction, functional annotation and protein/peptide classification. Its ability to learn both local and global context makes it a powerful tool for modern computational biology.

Recent advances in deep learning based multimodal peptide information include xBitterT5 [26]. It is a multimodal framework designed for bitter peptide identification that integrates BioT5+ transformer-based embeddings derived from peptide sequences with chemical representations. This combination enhances prediction accuracy, enabling researchers to identify specific amino acid regions contributing peptide bitterness. This framework has practical applications in food research, particularly in peptide design. Similarly, ABCP_finder is a deep learning-based tool developed for peptide prediction targeting to breast cancer. Like xBitterT5, it utilises the transformers-based embeddings to capture rich contextual, structural, and functional information from peptide sequences.

Despite of the recent advances, no computational tool has been developed to specifically identify ABCPs. Existing predictors can classify general ACPs and not tuned to separate ABCPs. In this study we present ABCP_finder, that performs classification of ABCPs and non-ABCPs using pretrained transformer embeddings-based framework. Overall, this framework enables high-throughput screening, significantly reducing workload and accelerating the identification of promising peptide candidates.

## 2. Methodology

The overall methodology is described below with data collection, homology-aware dataset Splitting, feature extraction, model architecture, training accuracy, performance evaluation and external validation. The architecture of ABCP_finder is provided in figure 1.

**Fig 1.**
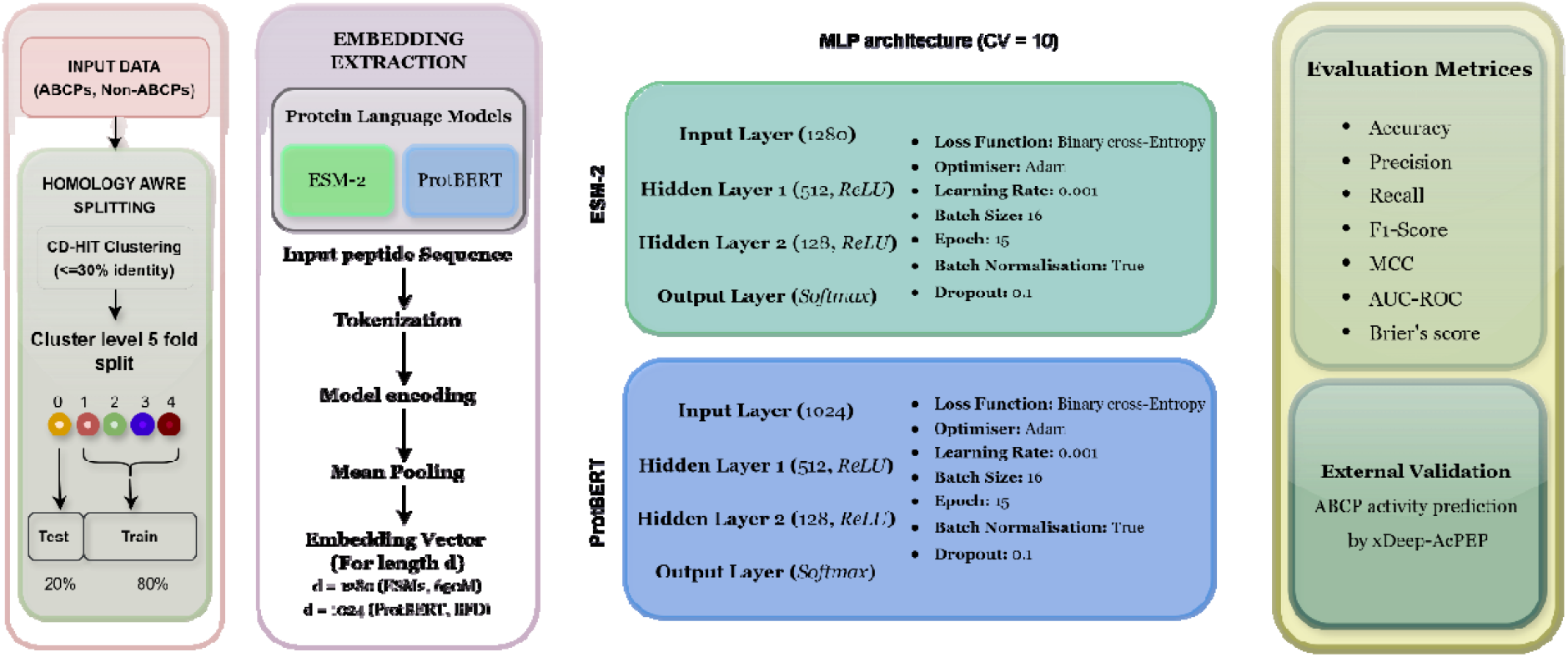
Overall architecture of ABCP_finder.

## a. Data Collection

ABCP_finder is built on carefully curated peptide data from well-established sources. ABCPs (positive dataset) were collected from the CancerPPD2 database [27], while non-ABCP (negative datasets) was obtained from the ACP-ML classifier [28], which is a recently developed and pioneering classifier trained on updated dataset of anti-cancer peptide (ACPs) and non-ACPs. To construct reliable negative dataset, we combined both general ACPs and non-ACPs, while excluding ABCPs.

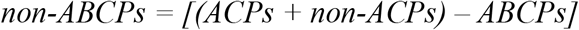

ABCPs and non-ABCPs were capped with the length less than 50 amino acids. Removal of duplicates and non-standard sequences (sequence contains non-standard amino acid) the final dataset comprises of 301 ABCPs and 1,052 non-ABCPs. This careful curation ensured that there was no overlap between positive and negative datasets. This helped in preventing data leakage and improving the robustness and reliability of the model during training and evaluation.

### b. Homology-Aware Dataset Splitting

To ensure a rigorous and unbiased evaluation, a homology aware dataset splitting strategy was adopted instead of conditional random splitting. Random splits might result in data leakage and overstated performance by placing similar or homologous sequences both in training and test dataset. To resolve this sequence redundancy was resolved by a clustering-based methodology. All peptide sequences were clustered utilizing the CD-HIT[29] method under rigorous similarity criteria, wherein sequences exhibiting ≥30% identity and ≥80% coverage of the shorter sequence were categorised into the same cluster. Sequences more than 30% identity may share structural and functional similarities.

CD-HIT clustering produced 5 non-redundant clusters from the combined dataset. After clustering the dataset was partitioned at the cluster level instead of individual sequence level. Each cluster was treated as an indivisible unit and randomly assigned to one of five mutually exclusive folds. This ensured that all the sequences within a cluster were confined to a single fold. Out of five folds, four folds (ABCP : non-ABCP :: 240 : 935) was used for training data and one-fold (ABCP : non-ABCP :: 61:117) was reserved for independent test set. This strategy guarantees that no sequence should share significant similarity, provides more realistic assessment for model performance on unseen test data. To validate the efficiency of homology-based splitting, pairwise sequence similarity between training and test dataset was evaluated. A sequence identity distribution analysis was generated to confirm the homology aware splitting is able to separate the independent test set from training set properly.

This technique increases the robustness and biological relevance of the evaluation framework by guaranteeing that the model is tested on evolutionarily distant sequences, thereby better emulating real-world prediction scenarios.

### c. Feature extraction strategy

Biological meaningful features were generated using state-of-the-art pretrained protein language models were used. These language models encode peptide sequences into dense numerical vectors. Specifically, ESM2 and ProtBERT, transformer-based protein language models pretrained on large scale protein sequence databases were used to extract embeddings from the positive and negative datasets. Both models were evaluated in this study because they represent different protein language model design philosophies. ProtBERT is an alignment informed pretrained model while ESM2 is alignment free and evolutionary contexed pretrained model. Hence, it remains unclear which representation suites better for capturing the short, functionally distinct peptides.

Sequences from both datasets were tokenized by inserting whitespace between individual amino acid (“QAY” → “Q A Y”) to match the input format required by both pretrained models. These tokenized sequences were then passed through both the transformers models to get contextualized hidden-state representations for each residue. These representations capture rich information related to sequence context, evolutionary patterns, and functional motifs. Mean pooling was chosen over CLS-token pooling because its aggregate information from every residue rather than relying on a single summary token. This results a single vector embedding per peptide. In this way, embeddings contain better information about short-range motif responsible for ABCP activity.

All the related information to these pretrained models are summarized below (Table 1). The generated feature vectors from ProtBERT and ESM2 were subsequently utilized as inputs for classification. In this way, the foundation of the proposed ABCP_finder framework is formed.

**Table 1.**
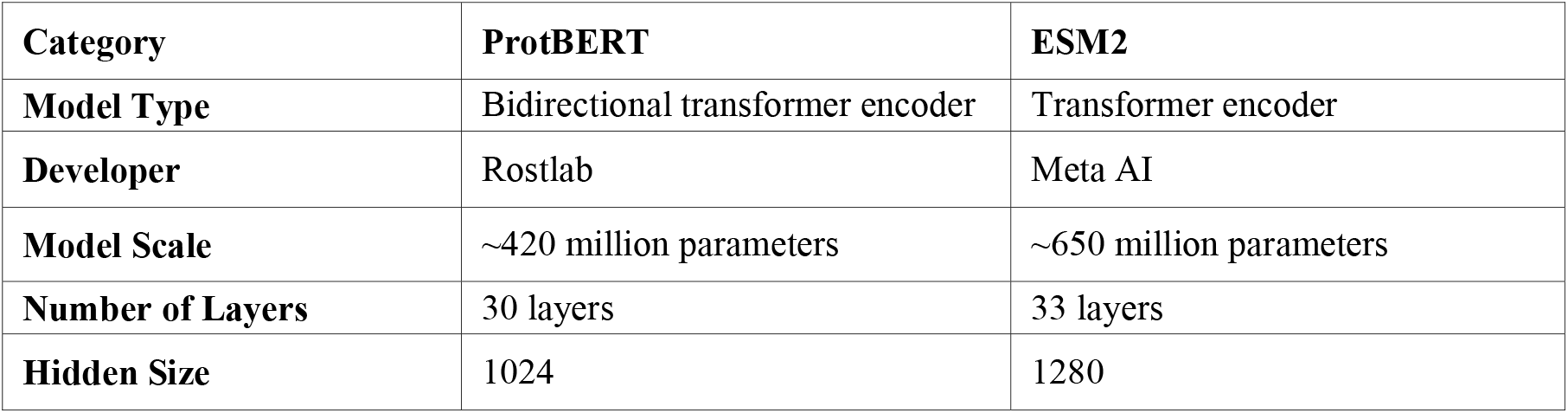
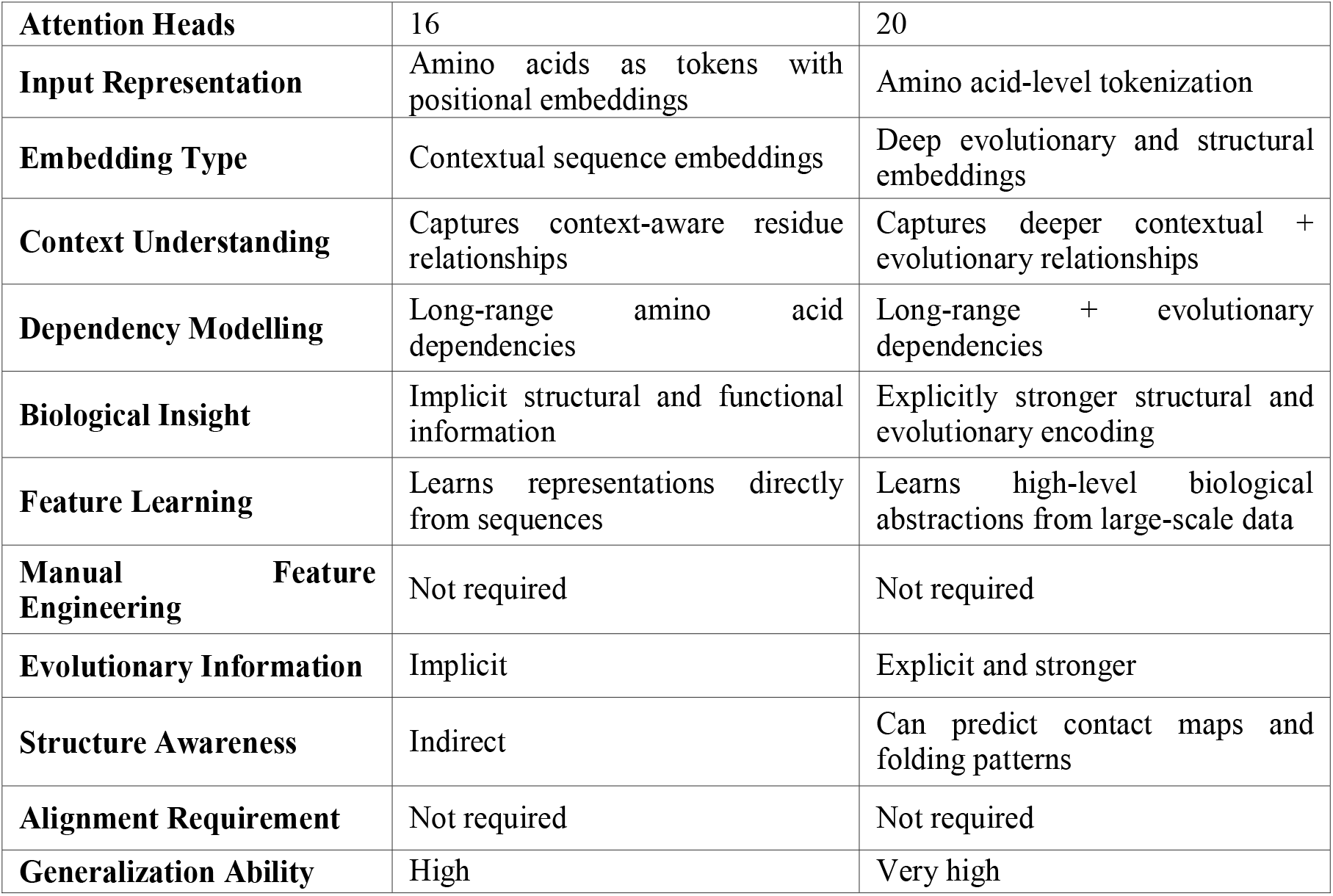
A comprehensive comparison of ProtBERT and ESM2 pretrained transformers.

### d. Model Architecture and Training Strategy

For the classification challenge, a multi-layer perceptron [30] was utilized. 10-fold grid search cross validation was used for hyperparameter training. This approach tested several parameter combinations to select the optimum hyperparameters. The tuned models are described below.

The input to the network is a fixed length embedding vector *x* ∈ *R*^*d*^ where *d* represents the dimensionality of the pretrained embedding, using a series of nonlinear processes, discover discriminative patterns between ABCPs and non-ABCPs.

In the first hidden layer, the input vector is linearly transformed using weights *W*_*1*_ and bias *b*_*1*_, followed by batch normalization [31] and a nonlinear activation:

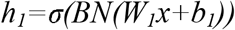

This stage enables the model to learn a collection of intermediate feature representations while stabilizing training with batch normalization, reducing internal covariate shift.

The second hidden layer further refines these learned representations:

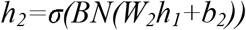

Here, the network captures higher-level abstractions by combining features from the previous layer, enabling it to model complex, nonlinear relationships within the embedding space.

The output layer then performs a final linear transformation:

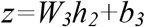

which produces raw prediction scores (logits) for each class. These logits are converted into probabilities using the *softmax* [32] function:

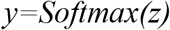

ensuring that the output values lie between 0 and 1 and sum to unity, making them interpretable as class probabilities.

The activation function used in the hidden layers is the Rectified Linear Unit (*ReLU*) [33], defined as:

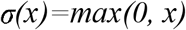

*ReLU* introduces nonlinearity into the model, allowing it to learn complex decision boundaries while maintaining computational efficiency.

*x* ∈ *R*^*d*^ denotes the input embedding vector of dimension d, derived from pretrained models, where each element represents learned sequence features. The matrices *W*_*1*_, *W*_*2*_, *W*_*3*_ correspond to the trainable weight parameters of the first hidden layer, second hidden layer, and output layer, respectively, while *b*_*1*_, *b2* and *b3* represent their associated bias vectors. The vectors *h*_*1*_ and *h*_*2*_ denote the intermediate hidden representations generated after the first and second layers, capturing progressively higher-level abstractions of the input features. The vector z represents the output logits, i.e., unnormalized scores for each class prior to probability conversion. The final prediction y corresponds to the class probability distribution obtained after applying the *softmax* function. The function σ(⋅) denotes the nonlinear activation function, specifically the Rectified Linear Unit (*ReLU*), which introduces nonlinearity into the model. The operator *BN(*⋅) represents batch normalization, which standardizes intermediate activations to stabilize and accelerate training.

To enhance generalization and reduce overfitting, dropout [34, 35] with a probability of *p=0*.*1* is applied after each hidden layer. This regularization technique randomly deactivates a fraction of neurons during training, preventing the network from becoming overly reliant on specific features and encouraging more robust feature learning.

The final layer consists of two output neurons corresponding to the binary classification task (ABCP vs non-ABCP), enabling the model to assign a probability score to each class.

The model parameters were optimized by *Adam* optimizer [35], which adapts learning rates for each parameter, based on first and second moment estimates of gradients. The optimization objective was to minimize the categorical cross-entropy loss which is suitable for binary classification. Training was performed with a learning rate of 0.001, with batch size of 16, and for a total of 15 epochs. The dataset was shuffled at each epoch to ensure the stochasticity and reduce bias during training.

During each iteration the model parameters were updated using back propagation:

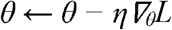

where θ represents model parameters and η is the learning rate.

This training methodology facilitates the fast learning of nonlinear decision limits over high-dimensional embedding spaces, allowing the model to successfully identify ABCPs from non-functional sequences.

### e. Performance Evaluation

The performance of the trained model was evaluated with the independent test dataset to ensure unbiased and complete assessment of the model. Diverse complementing metrics were utilized to assess various facets of classification performance, encompassing discrimination, dependability, and error distribution.

#### Accuracy

served as the principal criterion for assessing the overall correctness of predictions, defined as the ratio of correctly classified samples to the total number of samples:

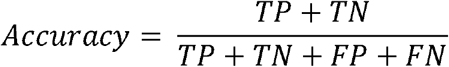

Where TP, TN, FP, FN denotes true positive, true negative, false positive and false negative, respectively. While accuracy provides general overview, it can be misleading in imbalanced dataset.

#### Precision

quantifies the ratio of predicted positives samples that are truly positive:

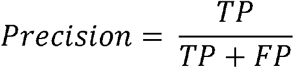

This metrics is particularly important in when the cost of false positives is very high, ensuring that predicted ABCPs should be reliable.

#### Recall (Sensitivity)

measures the ability of model to correctly identify true positive sample out of all positives:

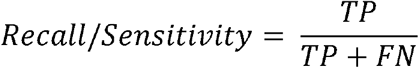

It reflects how model is capturing actual ABCPs and is critical in breast cancer applications where missing true positive can be problematic.

#### Specificity

evaluates the model correctly identify true negative samples out of all negatives:

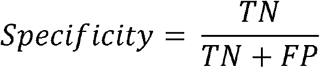

This metrics ensures that non-ABCPs are not incorrectly classified as ABCPs.

#### F1-score

represents the harmonic mean of precision and recall:

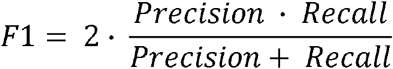

It provides a balance measure when both the false positive and false negatives are important.

#### Matthew’s correlation coefficient (MCC)

is a robust and important metrics that consider all elements of confusion metrics and particularly used in imbalanced dataset [36].

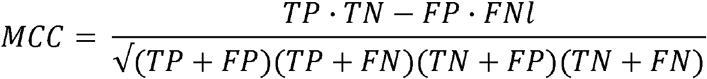

MCC ranges from -1 to +1, where +1 indicates the perfect prediction, 0 indicates random prediction, and -1 represents total discordance.

#### Area Under the Receiver Operating Curve (AUC-ROC)

evaluates the model’s ability to distinguish between ABCP and non-ABCP across different classification threshold. A higher AUC indicates better separability between ABCP and non-ABCP.

#### Brier score (BS)

[37] measures the accuracy of probabilistic predictions and reflects model calibration:

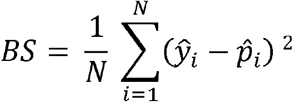

Where N is the total number if samples, *ŷ* ∈ [1] represent true class label, and and 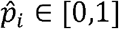 denotes the predicted probability of a sequence being an ABCP. Lower values illustrate better calibrated predictions.

Additional evaluation analysis like **Calibration curve** and **probability distribution** also performed. Calibration curve compares prediction probability with observed outcomes; this is particularly important in decision making in biomedical applications. Probability distribution of predicted probability for ABCP and non-ABCP. It shows the degree of separation of classes and effectiveness of decision threshold.

### f. Validation with xDeep-AcPEP

Random unclassified peptide sequences were retrieved from the PeptideAtlas database.ABCP_finder was utilized to predict the classes of each peptide. High-quality predictions with a probability greater than 0.70 were filtered. All selected peptides were subsequently evaluated using xDeep-AcPEP [16]. It is a deep learning framework that contains a module for predicting ACP activity against breast cancer. Spread, central tendency, and variability of the anticipated IC□□ values were analysed. This offers an intuitive representation of the activity range and facilitating comparison amongst peptides.

## 3. Results and Discussion

### a. Sequence Identity Distribution

The pairwise sequence identity distribution between training and testing dataset was computed and visualized. The result shows that the majority of the sequence pairs exhibit low similarity, predominantly within 0-20% identity range. Only a small fraction is approaching to the threshold of 30% (figure 2). This confirmed homology based splitting strategy successfully minimized the presence of homologous sequence across the training and testing data.

**Fig. 2.**
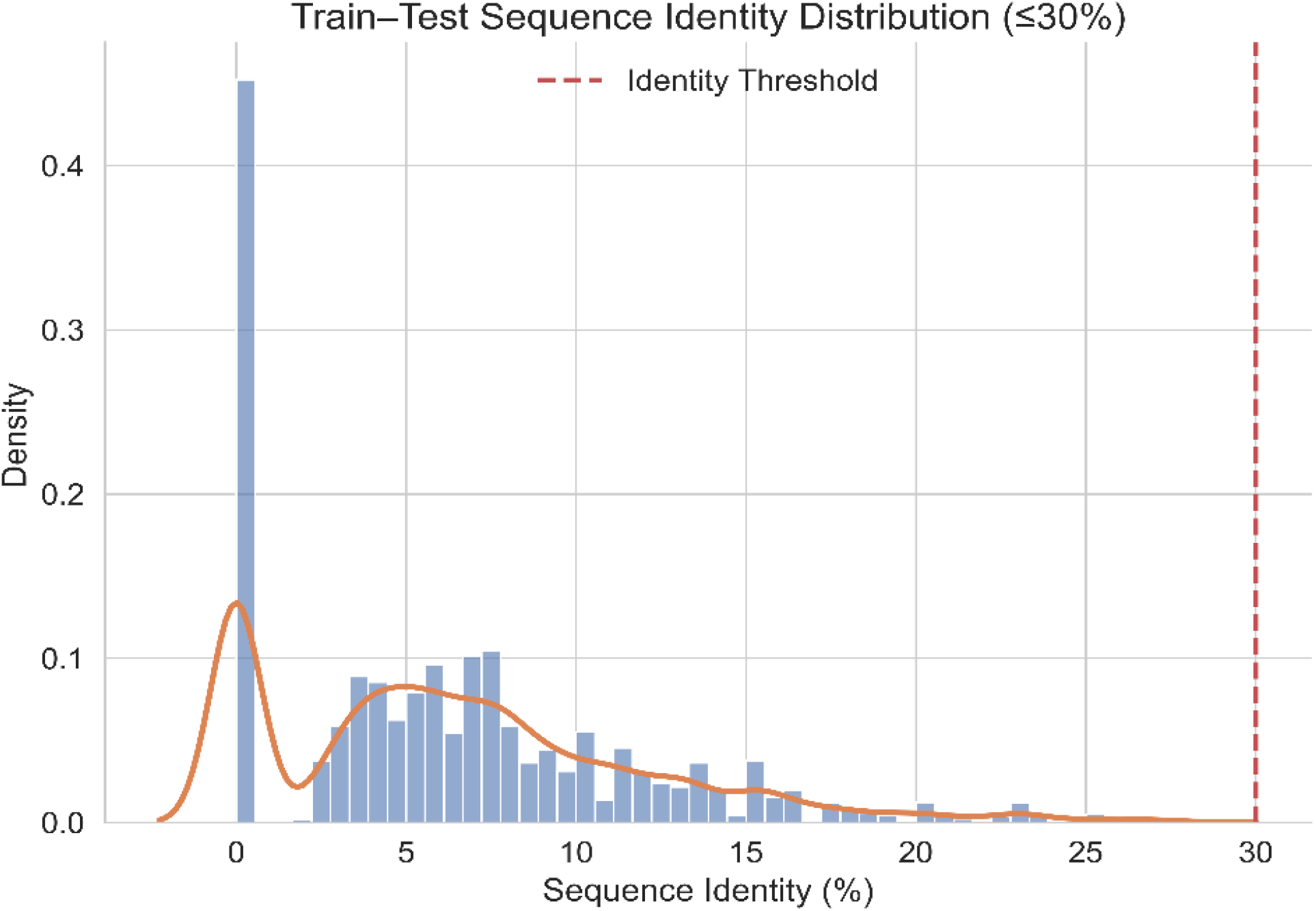
Plot illustrating train-test sequence identity distribution, confirming effective homology-based splitting with minimal sequence similarity.

### b. Model Performance

The model’s performance was evaluated using the test dataset in an imbalance setting, where ABCPs as the minority class relative to non-ABCPs. In such scenarios, accuracy is not only sufficient to evaluate model effectiveness; therefore precision, recall, F1-score, MCC, AUC, and BS were considered to evaluate the model shown in figure 3.

**Fig. 3.**
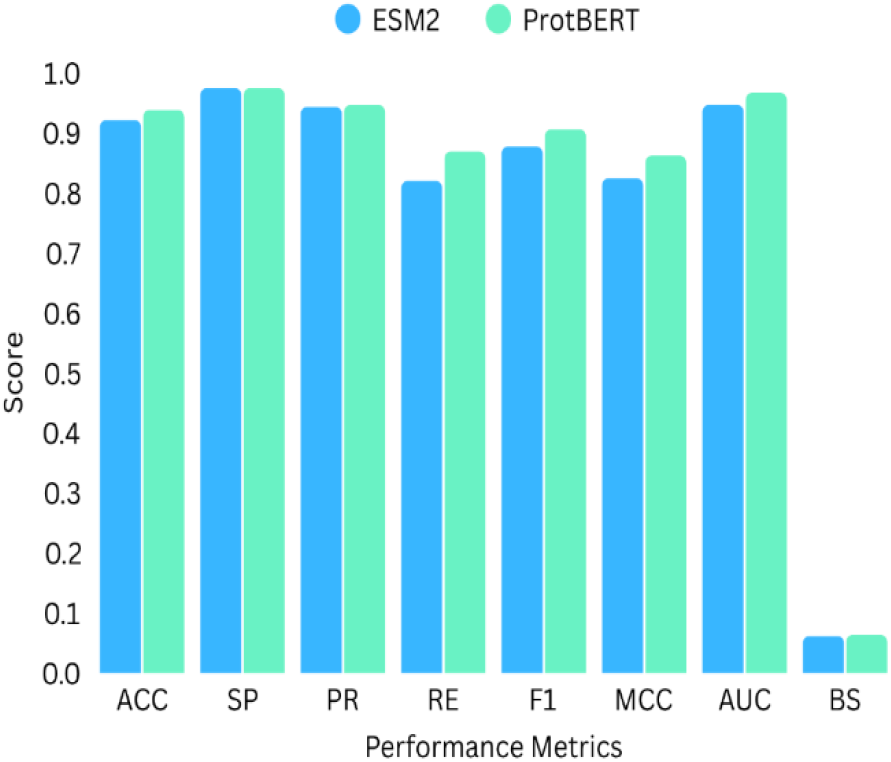
Evaluation metrices for ProtBERT and ESM2 based models, where ACC is accuracy, SP is specificity, PR is precision, RE is recall, F1 is F1-score, MCC is matthew’s correlation coefficient and BS is Brier score.

The ESM2-based model achieved an accuracy of 92.13%, with specificity of 97.43%, precision of 94.33%, recall of 81.96%, F1-score of 87.71%, MCC of 0.8241, AUC of 94.66% and BS of 0.0613. In comparison ProtBERT-based model shown slightly higher performance, achieving an accuracy of 93.82%, specificity of 97.43%, precision of 94.64%, recall of 86.88%, F1-score of 90.59% and MCC of 0.8618, AUC of 96.67% and the BS of 0.0633.

Both models showed excellent specificity (97%), which indicates a strong ability to correct identification of non-ABCPs. However, ProtBERT achieved the high recall compared to ESM2 based model (86.88% vs 81.96%) suggesting improved sensitivity in detecting true ABCPs. F1-score (90.59% vs 87.71%) and MCC (0.8618 vs 0.8241) is also higher for ProtBERT based model compared to ESM2 based model, which indicates the better balance between precision and recall.

The AUC of ProtBERT-based model is also higher than that of ESM2 based model (96.67% vs 94.66%), which demonstrate the good discrimination capability. Although ESM2-based model has achieved slightly better BS (0.0613 vs 0.0633) indicating marginally better probability calibration, but the difference is very minimal. ProtBERT-based model consistently outperformed ESM2-based model across all critical metrices like recall, F1-score, MCC and AUC for imbalanced classification problem.

The result demonstrate that both have achieved strong overall performance, however ProtBERT-based model consistently outperformed ESM2-based model to identify ABCPs with high recall, F1-score, MCC and AUC. This makes ProtBERT-based model more suitable for practical applications, for accurate detection of ABCP is critical. Although ESM2-based model is providing slightly better calibration as reflected as reflected by a lower BS, but overall predictive performance supports ProtBERT-based model. The improved sensitivity and balances performance of ProtBERT shows its effectiveness in handling imbalanced biological dataset. The stronger performance of the ProtBERT-based rather than ESM2-based model, may reflect differences in how the two language models represent short, functionally specialised peptide sequences. ESM2 embeddings provide a higher-capacity representation and not able to contextualise well on short ABCPs. Furthermore, given the relatively small and imbalanced dataset, the classifier may benefit from ProtBERT’s more compact embedding space (1024 dimensions) compared with the higher-dimensional ESM2 representation (1280 dimensions).

These findings suggest that pretrains transformer models-based embeddings, particularly from ProtBERT provide a robust framework for ABCP prediction. This can significantly aid in the discovery and screening of potential ABCPs.

### c. Confusion metrics, ROC curve, Training curves, Calibration curve

The confusion metrics provides a detailed breakdown of classification outcomes. For ESM2-based model and ProtBERT-based models, (114,114) non-ABCPs are correctly identified (True Negative), (3,3) non-ABCPs are incorrectly identified as ABCPs (False Positive), (8,11) ABCPs are incorrectly predicted as non-ABCPs (False negative), and (53,50) ABCPs are correctly predicted (True Positive) respectively. The model rarely misclassifies both the classes and maintains a strong recall for successful identification of ABCPs. Since the number of false negatives is relatively very low than the imbalance, indicating that model is very effective in predicting the critical minority class in prediction.

ROC curve provides trade-off between true positive rate and false positive rates across different classification thresholds. In both ESM2-based model and ProtBERT-based models, the curve is close to the top-left corner, which shows the near to ideal scenario of high sensitivity and low false positive rate. This result confirms that both the models can effectively distinguish between ABCPs and non-ABCPs sequences across wide range of threshold. Based on the AUC ProtBERT-based model is found to be better than ESM2-based model in terms of discriminative capability in imbalanced dataset, as it reflects the robustness of the model independent of class distribution.

Training curves illustrates the learning behaviour of the model over every epoch. Loss and accuracy for ProtBERT-based model and ESM-based model both are showing similar trend. Loss curve steadily decreases and accuracy curve consistently increases, this smooth and monotonic curve indicates the stable learning process, effective optimization with no sign of overfitting. There is no divergence between loss and accuracy curve, no sudden fluctuation suggesting model has been generalised well for unseen data.

The calibration curve assesses the alignment between projected probability and actual observed outcomes indicating that predicted probabilities are meaningful and trustworthy. Both the models showed reasonably good calibration with predicted probabilities. This indicates that the model provide reliable probability estimate. However, some deviation has been observed in both the models in mid probability regions, suggesting slight over-and-under confidence in prediction. A threshold (>=0.7) for both the models will be the optimum threshold for both the models to select the predicted ABCPs for experimental validation. In this case, ABCPs with strong predicted likelihoods are considered to reduce false positives. This is further supported by low BS (0.06) of both the models shows that probabilistic predictions are correct and helps in risk awareness decision making. The confusion metrics, ROC curve, training curves, calibration curve and prediction probability threshold is provided in figure 4.

**Figure 4.**
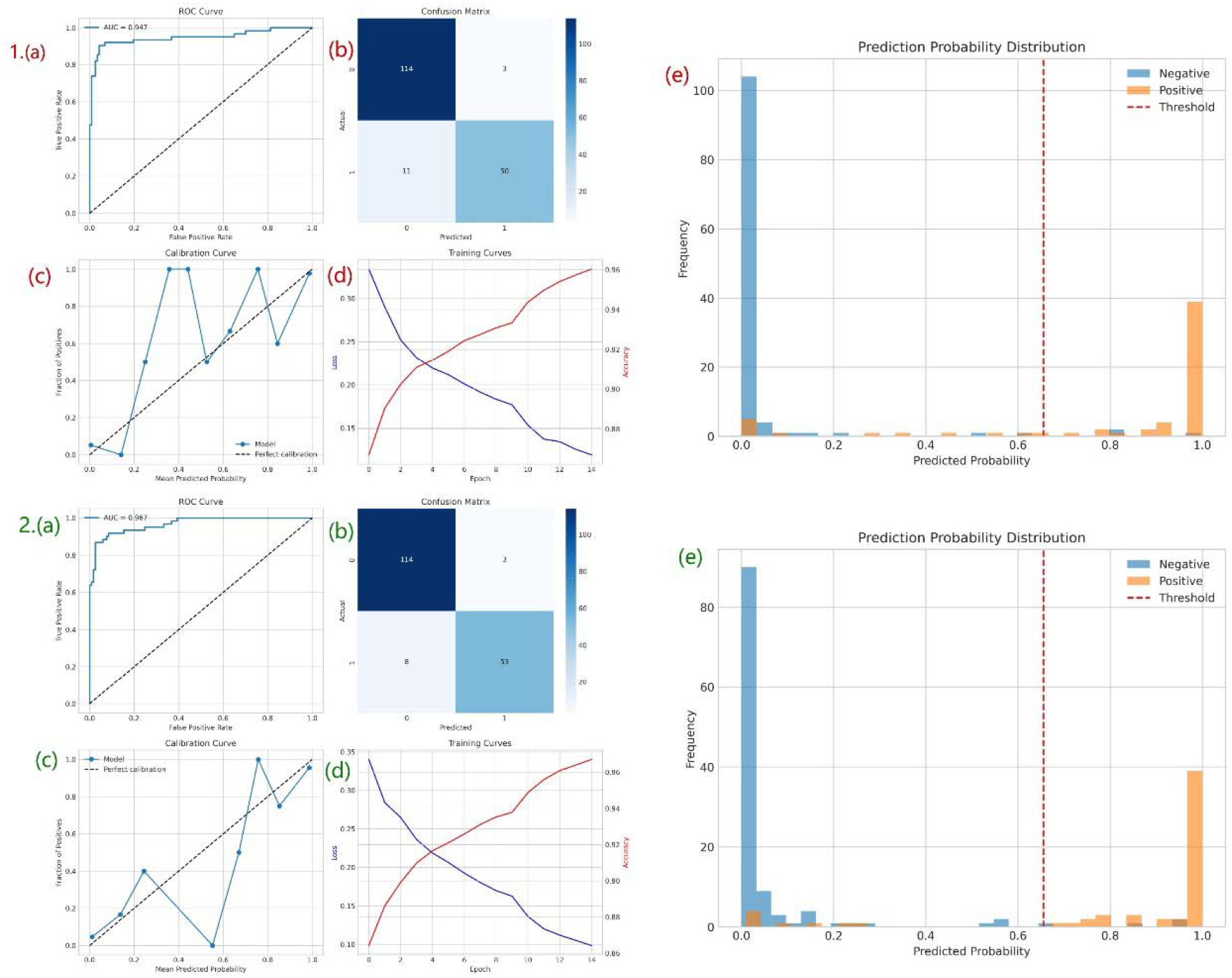
Performance evaluation of transformer-based models for anti-breast cancer peptide (ABCP) classification using ESM2 and ProtBERT embeddings. The upper panel (1) shows the results of the ESM2-based model, while the lower panel (2) presents the performance of the ProtBERT-based model. For both models, (a) the ROC curve illustrates the overall classification ability through the Area Under the Curve (AUC); (b) the confusion matrix shows the correctly and incorrectly classified positive and negative samples; (c) the calibration curve reflects how closely the predicted probabilities match the actual outcomes; (d) the training curves represent changes in loss and accuracy over different training epochs; and (e) the prediction probability distribution shows how well the model separates positive and negative classes, along with the selected decision threshold marked by the red dashed line.

### d. Final Validation

To further assess the potential activity of the peptides identified by ABCP_finder, 256,889 (out of 3,022,830) high-confidence peptides with a prediction probability >0.70 were selected for further analysis using xDeep-AcPEP [16]. The predicted IC□□ values were visualized using a violin plot (Figure 5).

**Figure 5.**
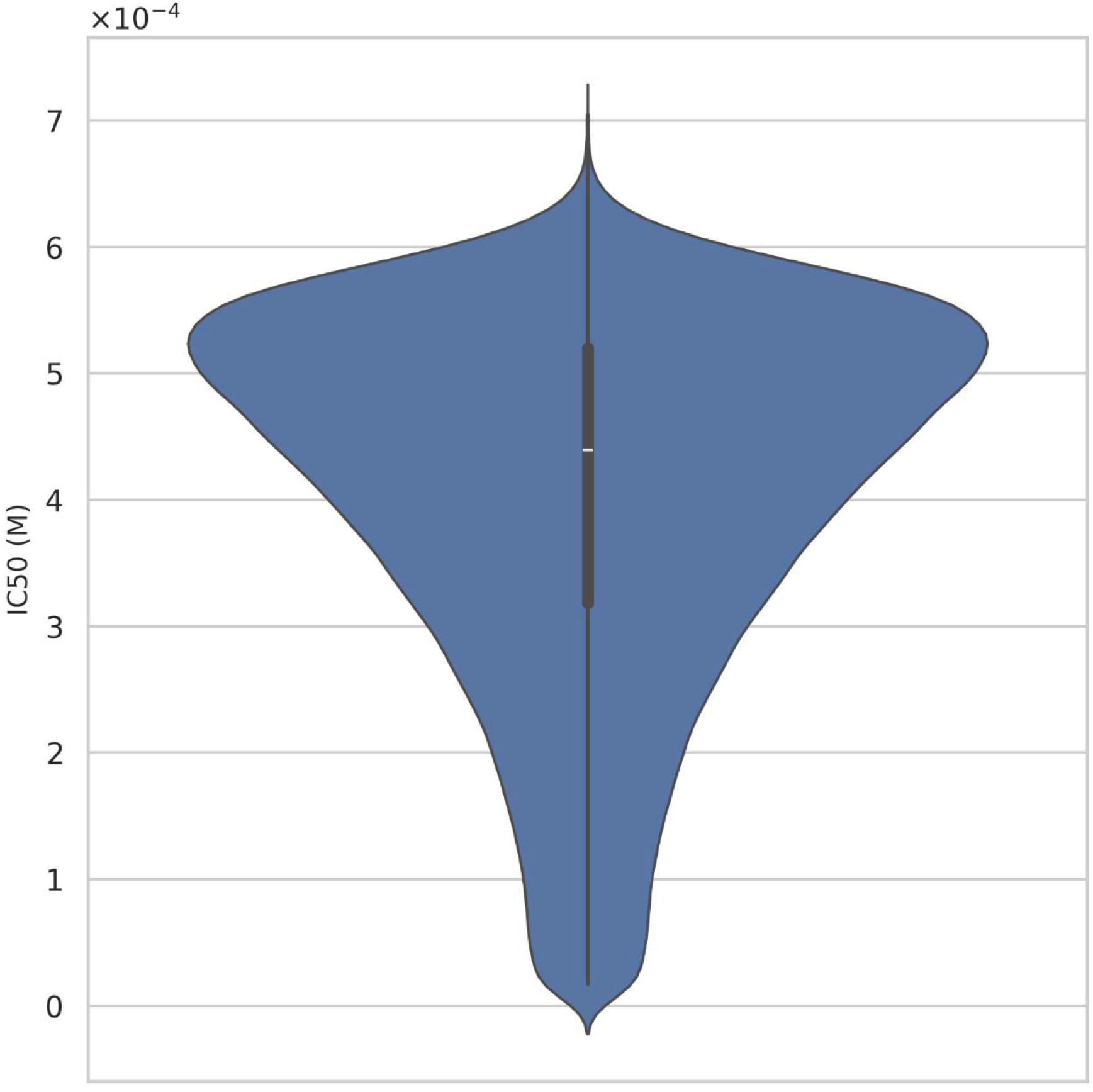
Violin plot illustrating the unknown peptides predicted from ABCP_finder as ABCP and their IC□□ values predicted from xDeep-AcPEP tool.

The distribution shows that most predicted IC□□ values fall within a relatively low micromolar-to-submillimolar molar range, mainly between approximately 1 × 10□□ M and 6 × 10□□ M. The median IC□□ is around 4.4 × 10□□ M. A large proportion of the peptides are concentrated around this region, as indicated by the wider part of the violin, while the overall distribution remained within the predefined concentration threshold used for ABCP activity assessment in the xDeep-AcPEP. Importantly, the distribution indicates that the peptides classified as ABCPs by ABCP_finder are associated with favourable predicted IC□□ values rather than being randomly distributed across the concentration range.

The concentration distribution therefore provides an independent computational assessment supporting the biological relevance of the ABCP_finder predictions. The agreement between ABCP classification and favourable predicted IC□□ values suggests that the model is capable of enriching for peptides with potentially stronger anticancer activity.

### e. Biological Implications

ABCP_finder has important biological significance in the discovery of ABCPs and supports precision therapeutics by identified potential ABCPs directly from the sequence information. Traditional methods that heavily dependent on manual feature engineering and sequence alignment, whereas ABCP_finder used transformer-based protein language models to learn deep biological information automatically from the peptide sequences. This allows model to capture properties such as sequence motifs, structural patterns, evolutionary relationships, amphipathicity, etc, which are strongly associated to its activity.

A major strength of this model is its ability to perform well under imbalanced conditions, where true ABCPs are much fewer than non-ABCPs. This is biologically important rare functional peptides are often the most valuable candidates. The higher recall and F1-score of the ProtBERT-based model indicate better detection of rare but therapeutically relevant to reduce the chance of missing strong candidates. The use of probability-based prediction further improves practical usefulness by allowing researchers to prioritize only high confidence peptides for validation. External validation showed that the most predicted ABCPs has favourable IC□□ values that confirms real biological relevance to make ABCP_finder a reliable tool for ABCP discovery.

ProtBERT and ESM2 embeddings are learned directly from large-scale protein sequence data without explicit feature engineering. These representations may implicitly capture biologically relevant properties, such as sequence motifs, secondary-structure propensity, and amphipathicity. However, this study did not directly investigate which features were used by the classifier for prediction. Future interpretability analyses, including attention-weight visualization and embedding-space feature attribution, could help identify the sequence characteristics underlying ABCP prediction.

## 4. Conclusion

In this study, we developed ABCP_finder, the first dedicated computational tool specifically designed for ABCPs classification. ABCPs and non-ABCPs were taken from CancerPPD2 and ACP-ML. To make classification more realistic, train-test split was performed with homology awareness with CD-HIT at 30% identity and 80% coverage. This helped prevent similar sequence appearing in both train and test dataset, reducing the data leakage and ensure more reliable evaluation.

Among the tested transformer-based models, ProtBERT performed better than ESM2, specially in identifying the minority ABCP class. It achieved accuracy of 93.82%, recall of 86.88%, F1-score of 90.59%, MCC of 0.86 and 96.67% AUC, which shows strong predictive performance even in under imbalanced condition. The low BS indicates that model is well calibrated and reliable for decision making. Based on the calibration analysis, a probability cutoff of 0.70 was selected as optimal threshold to identify high confidence ABCPs. These metrices allows better prioritization of promising ABCP candidate while reducing false positive prediction during downstream experimental validation.

Further validation with using xDeep-AcPEP showed that predicted ABCPs are falling as favourable IC□□ values, simulating real biological activity against breast cancer. Overall, ABCP_finder provides a reliable and practical platform for novel ABCP discovery and can accelerate future therapeutic development.

## 5. Limitations, and Future Directions

While ABCP_finder shows strong predictive ability and offers a fast computational way to screen anti-breast cancer peptides, there are some important limitations to keep in mind. The model was trained on relatively small and imbalanced dataset, which can affect how well it works on new or structurally different peptides. Furthermore, because the majority of the data originates from trials on specific breast cancer cell lines, the model may be learning cell line-specific patterns rather than universally applicable across all breast cancer subtypes. Currently, ABCP_finder conducts binary classification without providing information on potency, mechanism of action, or subtype specificity. Although, ABCP_finder is a valuable *in-silico* screening tool but the predictions must be experimentally validated before they are considered for therapeutic use. Another point to consider is that the model is entirely dependent on ProtBERT and ESM2 embeddings. Limitations or biases in those models may carry over into the predictions.

Looking ahead, there are several ways to improve the framework. Expanding the dataset with peptides evaluated across numerous breast cancer subtypes. More experimental data would assist in increasing robustness and generalizability. Incorporating quantitative activity data could allow the model to move beyond classification and would predict the IC50 values using transformers. Adding structural knowledge would also make the prediction more physiologically meaningful. Moreover, there is also space to integrate explainable AI methods to better understand which sequence feature are driving the prediction, it can boost the trust in the model’s output. Finally, developing a system that continuously updates the model with new experimental data will help reduce bias over time and make ABCP_finder as a strong tool for guiding ABCP discovery.

## 6. Tool and Data Availability

The supporting data and command line tool are available on GitHub at the following link: https://github.com/darkest-opson/abcp_finder_module and https://github.com/darkest-opson/ABCP_finder/. The ABCP_finder is both CPU and GPU supported to accelerate the drug discovery.

## 7. Funding availability

No Funding is utilized in the development of this tool.

## Notes

https://github.com/darkest-opson/abcp_finder_module

https://github.com/darkest-opson/ABCP_finder/

### Competing Interest Statement

The authors have declared no competing interest.

